# Caffeine extends lifespan by enhancing lysosomal lipolysis in *Caenorhabditis elegans*

**DOI:** 10.64898/2026.01.08.698302

**Authors:** Hyemin Min, Eunseok Kang, Gee-Yoon Lee, Laura Bahr, Arjumand Ghazi, Seung-Jae V. Lee

## Abstract

Caffeine is a globally consumed stimulant that has beneficial effects on biological processes including metabolism and aging, but its causal role in physiology remains incompletely understood. By using the roundworm *Caenorhabditis elegans*, here we show that caffeine extends lifespan by eliciting transcriptional remodeling that enhances lysosomal lipolysis. We found that transcriptomes of aged, caffeine-fed animals shifted toward youthful states. By comparing with three longevity-promoting regimens, including reduced insulin/insulin-like growth factor 1 (IGF-1) signaling, mild reductions in mitochondrial function, and dietary restriction (DR), we showed that caffeine induced a DR-like transcriptional program. Comparison with *eat-2* mutants (a genetic DR model) identified lysosomal lipases *lipl-1* and *lipl-2* as commonly upregulated genes. The induction of *lipl-1* and *lipl-2* was required for increased lifespan and reduced neutral lipid accumulation by caffeine intake. Together, these findings indicate that caffeine promotes longevity in a DR-like metabolic reprogramming by enhancing lysosome-driven lipolysis.

## INTRODUCTION

Caffeine, a key component of beverages such as coffee and tea, enhances arousal, vigilance, and psychomotor performance (Nehlig et al., 1992; Smith, 2002; McLellan et al., 2016). Moderate caffeine consumption is linked to reduced risks of various chronic diseases, including non-alcoholic fatty liver disease, cardiovascular diseases, Alzheimer disease and type 2 diabetes risk (Poole et al., 2017; Carlstrom and Larsson, 2018; Hayat et al., 2021; Gill et al., 2024). Caffeine intake is also implicated in reductions in body weight, body mass index, and waist circumference (Phung et al., 2010; Tabrizi et al., 2019). Moreover, caffeine extends lifespan in several model organisms, including the budding yeast *Saccharomyces cerevisiae* and the nematode *Caenorhabditis elegans* (Wanke et al., 2008; Sutphin et al., 2012; Du et al., 2018; Min et al., 2021). In *C. elegans*, caffeine (1–10 mM) extends lifespan and improves healthspan, with genetic data implicating the roles of stress response and insulin/insulin-like growth factor 1 (IGF-1) signaling (Sutphin et al., 2012; Du et al., 2018; Min et al., 2021). Despite these beneficial effects of caffeine on health, underlying molecular mechanisms, in particular, for organismal longevity remain poorly understood.

## MATERIALS AND METHODS

### C. elegans Strains

*C. elegans* strains were maintained at 20°C on standard nematode growth medium (NGM) plates seeded with OP50 (Park et al., 2025), unless stated otherwise. Detailed information of the strains used in this study is described in the Supplementary Information.

### RNA sequencing library preparation and analysis

Library preparation was performed using TruSeq Stranded mRNA Library Prep Kit, and sequencing was performed on the Illumina NovaSeq platform with paired-end reads. RNA sequencing data were analyzed following previous studies with minor modifications (Ham et al., 2022; 2024; Lee et al., 2024a; Lee et al., 2024b; Sohn et al., 2025). Detailed analysis methods are described in the Supplementary Information.

### Lifespan assays using *C. elegans*

Lifespan assays were performed as previously described (Ji et al., 2024; Lee et al., 2025a; Kwon and Lee, 2025) with minor modifications. Details are described in the Supplementary Information. Survival analyses were performed using the Online Application of Survival Analysis 2 (OASIS2, http://sbi.postech.ac.kr/oasis2) and OASIS portable (OASISp) (Han et al., 2016; Han et al., 2024).

### Microscopy and image analysis

Animals were mounted on a 2% agarose pad and anesthetized with 100□mM sodium azide (Sigma-Aldrich, 71289-5□G) prior to imaging (Jung et al., 2021; Kim et al., 2024; Min et al., 2025). Images were acquired using a Leica DFC9000 GT deep-cooled sCMOS camera and Leica LAS X imaging software (Leica Microsystems, Germany). Further details are provided in the Supplementary Information.

### Measurement of pharyngeal pumping rates

Pharyngeal pumping (feeding) of *C. elegans* was measured as previously described (Hwang et al., 2025; Lee et al., 2025b) with minor modifications. The pharyngeal pumping rate was counted for 30 seconds and subsequently converted to pumps per minute. Further details are described in the Supplementary Information.

### RNA extraction and quantitative RT-PCR (RT-qPCR)

RNA extraction and RT-qPCR were performed as previously described (Kim et al., 2023; Bong et al., 2024) with minor modifications. Further details are provided in the Supplementary Information. Primer sequences used for RT-qPCR analyses in this study are listed in the Table S1.

### Oil Red O staining

Oil Red O staining was performed as previously described (Lee et al., 2019) with minor modifications. See Supplementary Information for detailed methods.

## RESULTS

### Caffeine extends lifespan and elicits DR-like transcriptomic changes

We sought to investigate how caffeine intake affected lifespan and healthy aging in *C. elegans* and first confirmed the longevity-promoting effect of caffeine (10 mM) (Fig. 1a; Sutphin et al., 2012; Du et al., 2018; Min et al., 2021). We then performed RNA sequencing (RNA-seq) analysis of caffeine-fed animals (Fig. S1a) and compared the transcriptomic profile with those of young (day 1 of adulthood) and old (day 9 of adulthood) wild-type (WT) animals (Lee et al., 2021). We found that caffeine reversed the transcriptomic state of old animals to that of young animals (Fig. 1b, c). In addition, transcriptomic changes caused by caffeine and aging negatively correlated (correlation coefficient *r* = −0.76; Fig. S1b). By using Transcriptomic CLassification via Adaptive learning of Signature States (T-CLASS) (Lee et al., 2025c), we determined whether longevity conferred by caffeine feeding was similar to any of the three established longevity paradigms in *C. elegans*: dietary restriction (DR), reduced insulin/IGF-1 signaling (rIIS), and reduced mitochondrial function (rMF) (Bong et al., 2025). We found that the effect of caffeine feeding was most similar to that of DR (Fig. 1d). In addition, transcriptomic changes caused by caffeine supplementation and *eat-2* mutations, an established genetic DR model (Kapahi et al., 2017), positively correlated (correlation coefficient *r* = 0.8; Fig. 1e). Consistently, caffeine did not further increase the long lifespan caused by *eat-2* mutations (Fig. 1f), without affecting feeding rates (Fig. 1g). In contrast, caffeine further increased the long lifespan of nutrient-sensing defective sensory *osm-5* mutants (Fig. 1h), which display longevity (Apfeld and Kenyon, 1999). Thus, caffeine appears to promote longevity through mimicking DR, without reducing food intake or impairing nutrient sensing.

**Figure 1.**
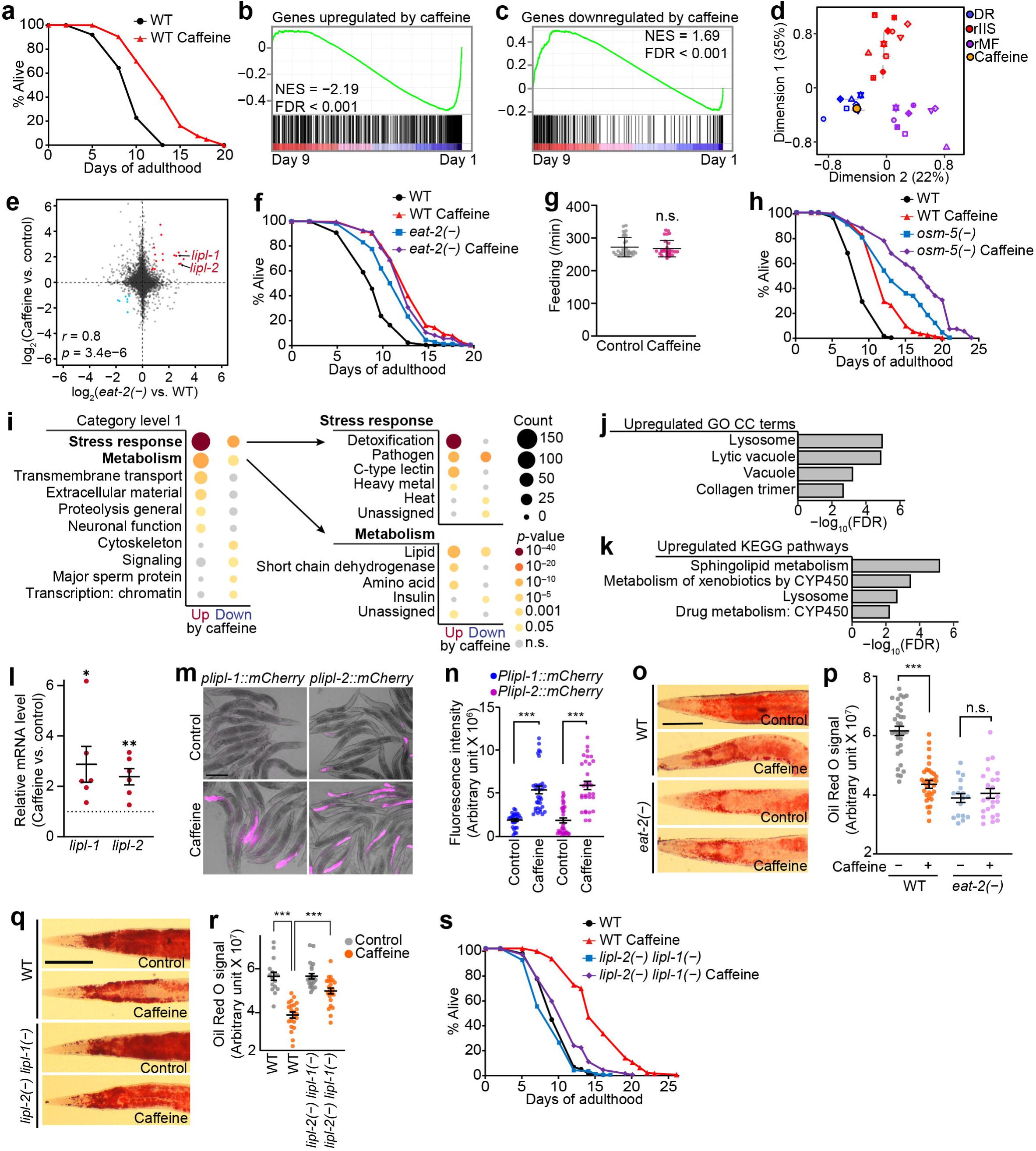
Induction of lysosomal lipolysis by caffeine feeding mediates dietary restriction (DR)-like longevity in *Caenorhabditis elegans*. (**a**) Lifespan extension by caffeine (10 mM) feeding in wild-type (WT) *C. elegans*. The same data sets are also shown in Fig. 1f. (**b**–**c**) Gene set enrichment analysis (GSEA) plots showing enrichment of genes up (b) or downregulated (c) by caffeine (10 mM) feeding during aging. NES, normalized enrichment score. FDR, false discovery rate. (**d**) Classification of transcriptomic changes caused by caffeine feeding using Transcriptomic CLassification via Adaptive learning of Signature States (T-CLASS) (Lee et al., 2025). (**e**) A scatter plot comparing gene expression changes induced by caffeine (10 mM) and by the *eat-2* mutations. *r*, Pearson correlation coefficient. See Table S2 for the list of genes commonly upregulated or downregulated in caffeine-fed animals and in *eat-2(ad1116)* [*eat-2(−)*] mutants. (**f**) Lifespan of WT and *eat-2(−)* mutants fed with or without caffeine. To visualize the differential effects of the caffeine on lifespan of WT and *eat-2(−)* mutants, the same WT data sets are also shown in Fig. 1a. (**g**) Pharyngeal pumping (feeding) rates of WT animals treated with caffeine (10 mM) compared with control (N = 30 for each condition, two independent trials). Error bars represent standard error of the mean (SEM). n.s., not significant. (**h**) The lifespan of WT and *osm-5(p813)* [*osm-5(−)*] mutants fed with or without caffeine. (**i**) WormCat terms overrepresented among genes upregulated or downregulated by caffeine at category levels 1 and 2. “Stress response” and “Metabolism” were the two most highly enriched terms (bold) among the upregulated genes at category level 1. *p*-values were calculated by using Fisher’s exact test. (**j**–**k**) Overrepresented gene ontology (GO) cellular component (CC) (j) and Kyoto Encyclopedia of Genes and Genomes (KEGG) pathway (k) terms significantly enriched among genes upregulated by caffeine. FDR, false discovery rate. (**l**) The mRNA levels of *lipl-1* and *lipl-2* were measured using RT-qPCR in WT animals fed with caffeine (n = 6). Dotted line indicates 1. Error bars represent SEM. *p*-values were calculated by using two-tailed Student’s *t*-test (\**p* < 0.05, \*\**p* < 0.01). (**m**–**n**) Expression of *lipl-1* and *lipl-2* monitored using the transcriptional reporter *plipl-1::mCherry* and *plipl-2::mCherry* animals fed with caffeine (N ≥ 27 for each condition, two independent trials). Scale bar, 200 μm. Error bars represent SEM. *p*-values were calculated by using a two-tailed Student’s *t*-test (\*\*\**p* < 0.001). (**o**–**p**) Representative images (o) and quantification (p) of Oil Red O staining in WT and *eat-2(−)* mutants fed with or without caffeine (N ≥ 17 for each condition, two independent trials). Scale bar, 75 μm. Error bars represent SEM. *p*-values were calculated by using two-tailed Student’s *t*-test (\*\*\**p* < 0.001). (**q**–**r**) Representative images (q) and quantification (r) of Oil Red O staining in WT and *lipl-2(glm79) lipl-1(glm77)* [*lipl-2(−) lipl-1(−)*] mutants fed with or without caffeine (N ≥ 16 for each condition, two independent trials). To visualize the differential effects of caffeine on lipid storage in WT and *lipl-2(−) lipl-1(−)* mutants, the same WT data sets are also shown in Fig. S1d and Fig. S1e. Scale bar, 75 μm. Error bars represent SEM. *p*-values were calculated by using one-way ANOVA (\*\*\**p* < 0.001). (**s**) The lifespan of WT and *lipl-2(−) lipl-1(−)* mutants fed with or without caffeine. To visualize the effects of the *lipl-1* and *lipl-2* mutations on caffeine-mediated longevity, the same WT data sets are also shown in Fig. S1f and Fig. S1g. Lifespan results in this figure represent pooled data from two independent experiments; results of individual replicates are provided in Table S3.

### Caffeine promotes longevity and reduces lipid levels by inducing lysosomal lipases

We further analyzed individual genes differentially expressed by caffeine (fold change > 1.5 and adjusted *p*-value < 0.05; Fig. S1c). By performing pathway enrichment analysis using WormCat (Holdorf et al., 2020; Higgins et al., 2022) and KEGG pathways (Kanehisa and Goto, 2000), we found that genes responsible for stress response were the most highly enriched term among upregulated genes by caffeine (Fig. 1i), concordant with previous reports (Min et al., 2015; Al-Amin et al., 2016). The second most highly enriched term among the upregulated genes was ‘Metabolism’ (Fig. 1i). In addition, KEGG pathway and gene ontology (GO) cellular component (CC) terms related to lysosomal activity, including ‘lysosome’ and ‘lytic vacuole’, were significantly enriched among caffeine-responsive genes (Fig. 1j, k). We then compared the transcriptomic profile of caffeine-fed animals and that of *eat-2* mutants (Tabrez et al., 2017). We found that *lipl-1* and *lipl-2*, genes encoding lysosomal lipases that drive lipophagy-linked lipid breakdown (O’Rourke and Ruvkun, 2013; Murphy et al., 2019), were commonly upregulated by caffeine and by *eat-2* mutations (Fig. 1e; See Table S2 for the list of commonly up or downregulated genes). We validated the upregulation of *lipl-1* and *lipl-2* by caffeine feeding using RT-qPCR (Fig. 1l) and transcriptional reporter strains (Fig. 1m, n). By using Oil Red O staining, we showed that caffeine feeding reduced overall lipid levels in WT, while not further decreasing it in *eat-2* mutants (Fig. 1o, p). Importantly, we found that the reduced lipid levels by caffeine intake was partially but significantly suppressed by *lipl-1(−)* and *lipl-2(−)* double mutations (Fig. 1q, r; see Fig. S1d, e for *lipl-1(−)* and *lipl-2(−)* single mutants data). Thus, caffeine appears to decrease fat levels by enhancing lysosomal lipolysis via upregulating *lipl-1* and *lipl-2*. Lastly, we showed that *lipl-1(−)* and *lipl-2(−)* double mutations largely suppressed extended lifespan conferred by caffeine feeding (Fig. 1s; see Fig. S1f, g for *lipl-1(−)* and *lipl-2(−)* single mutants data). Altogether, our data demonstrate that caffeine extends lifespan in a DR-like manner by reducing lipid accumulation through inducing lysosomal lipases.

## DISCUSSION

### Caffeine appears to promote longevity by enhancing lipid catabolism in

#### lysosome

In this study, we defined a link between lifespan extension and a lipid catabolic program induced by caffeine. We showed that caffeine increases lysosomal lipolysis through a DR-like transcriptional program and that this lipid catabolic axis is required for the pro-longevity effects. In humans, lipase-dependent lipid metabolism, including reduced post-heparin lipoprotein lipase activity and slower triglyceride clearance, decreases with age, along with broader lipid-metabolic remodeling that impairs fat mobilization (Spitler and Davies, 2020). Lower hepatic lipase activity and disrupted endothelial and lysosomal lipases are associated with cardiometabolic disease, age-dependent cognitive decline, and progressive liver pathology (Santamarina-Fojo et al., 2004; Reiner et al., 2014; Yun et al., 2019). Thus, impairments in lipolysis and triglyceride metabolism may serve as functional indicators of biological aging. If our study holds true in mammals, a simple dietary intervention, caffeine intake, may counteract these deficits and prevent aging and age-related diseases (López-Otín et al., 2023; Kang et al., 2024; Park et al., 2024; Yan et al., 2024; Kim and Lee, 2025), by enhancing lysosomal lipolysis and triglyceride turnover. Future studies should determine whether this coupling also operates in mammals by assessing changes in lysosome-dependent lipolysis and functional aging markers upon caffeine consumption. In addition, a more detailed understanding of the upstream molecular mechanisms will be critical. Caffeine primarily functions as an antagonist of adenosine receptors, thereby inhibiting adenosine signaling and altering downstream pathways that regulate cellular metabolism (Ribeiro and Sebastiao, 2010; Jacobson et al., 2020). Identifying the transcription factors and characterizing adenosine receptor-dependent signaling cascades that drive caffeine-induced lipid remodeling will be essential for developing caffeine-based interventions that promote healthy aging.

## AUTHOR CONTRIBUTIONS

*Author contributions*: H.M., E.K., G.-Y.L., and S.-J.V.L. contributed to designing the research. H.M., E.K, and G.-Y.L. performed data analyses. L.B., and A.G. contributed key experimental materials. H.M., E.K., and G.-Y.L. performed data curation. H.M. and E.K. performed experiments. H.M., E.K., G.-Y.L., and S.-J.V.L. wrote the manuscript. All authors read and approved the manuscript.

## ACKNOWLEDGMENTS

We thank all the Lee lab members for comments on the manuscript and discussion. We appreciate Dr. Yong-Hee Shim for technical assistance with RNA-seq experiments and Seung-Chul J. Lee for assistance with RNA-seq data analysis.

## DECLARATION OF GENERATIVE AI AND AI-ASSISTED TECHNOLOGIES IN THE WRITING PROCESS

During the preparation of this work the authors used ChatGPT 5.1 (OpenAI) to improve language clarity of the manuscript. After using this tool, the authors reviewed and edited the content as needed and take full responsibility for the content of the publication.

## DATA AVAILABILITY STATEMENT

All raw and processed sequencing data has been submitted to the NCBI Gene Expression Omnibus (GEO; https://www.ncbi.nlm.nih.gov/geo/) under accession number GSE312248. All the other data that support the findings of this study are available from the corresponding author upon reasonable request.

## SUPPLEMENTARY MATERIALS

### Supplementary Information

**Table S1**. List of RT-qPCR primers used in this study

**Table S2**. List of genes commonly up or downregulated by caffeine and by *eat-2(−)* mutations

**Table S3**. Raw data and statistical analysis

## REFERENCES

1. Al-Amin, M., Kawasaki, I., Gong, J., and Shim, Y.H. (2016). Caffeine Induces the Stress Response and Up-Regulates Heat Shock Proteins in *Caenorhabditis elegans*. Mol Cells 39, 163–168.

2. Apfeld, J. and Kenyon, C. (1999). Regulation of lifespan by sensory perception in *Caenorhabditis elegans*. Nature 402, 804–809.

3. Bong, D., Kwon, H.C., and Lee, S.V. (2025). Multilayered regulation of longevity in *Caenorhabditis elegans*. Mol Cells, 100308.

4. Bong, D., Sohn, J., and Lee, S.V. (2024). Brief guide to RT-qPCR. Mol Cells 47, 100141.

5. Carlstrom, M. and Larsson, S.C. (2018). Coffee consumption and reduced risk of developing type 2 diabetes: a systematic review with meta-analysis. Nutr Rev 76, 395–417.

6. Du, X., Guan, Y., Huang, Q., Lv, M., He, X., Yan, L., Hayashi, S., Fang, C., Wang, X., and Sheng, J. (2018). Low Concentrations of Caffeine and Its Analogs Extend the Lifespan of *Caenorhabditis elegans* by Modulating IGF-1-Like Pathway. Front Aging Neurosci 10, 211.

7. Gill, H., Patel, N., Naik, N., Vala, L., Rana, R.K., Jain, S., Sirekulam, V., Jain, S.M., Khan, T., Kinthada, S., et al. (2024). An umbrella review of meta-analysis to understand the effect of coffee consumption and the relationship between stroke, cardiovascular heart disease, and dementia among its global users. J Family Med Prim Care 13, 4783–4796.

8. Ham, S., Kim, S.S., Park, S., Kim, E.J.E., Kwon, S., Park, H.H., Jung, Y., and Lee, S.V. (2022). Systematic transcriptome analysis associated with physiological and chronological aging in *Caenorhabditis elegans*. Genome Res 32, 2003–2014.

9. Ham, S., Kim, S.S., Park, S., Kwon, H.C., Ha, S.G., Bae, Y., Lee, G.Y., and Lee, S.V. (2024). Combinatorial transcriptomic and genetic dissection of insulin/IGF-1 signaling-regulated longevity in *Caenorhabditis elegans*. Aging Cell 23, e14151.

10. Han, S.K., Kwon, H.C., Yang, J.S., Kim, S., and Lee, S.V. (2024). OASIS portable: User-friendly offline suite for secure survival analysis. Mol Cells 47, 100011.

11. Han, S.K., Lee, D., Lee, H., Kim, D., Son, H.G., Yang, J.S., Lee, S.V., and Kim, S. (2016). OASIS 2: online application for survival analysis 2 with features for the analysis of maximal lifespan and healthspan in aging research. Oncotarget 7, 56147–56152.

12. Hayat, U., Siddiqui, A.A., Okut, H., Afroz, S., Tasleem, S., and Haris, A. (2021). The effect of coffee consumption on the non-alcoholic fatty liver disease and liver fibrosis: A meta-analysis of 11 epidemiological studies. Ann Hepatol 20, 100254.

13. Higgins, D.P., Weisman, C.M., Lui, D.S., D’Agostino, F.A., and Walker, A.K. (2022). Defining characteristics and conservation of poorly annotated genes in *Caenorhabditis elegans* using WormCat 2.0. Genetics 221.

14. Holdorf, A.D., Higgins, D.P., Hart, A.C., Boag, P.R., Pazour, G.J., Walhout, A.J.M., and Walker, A.K. (2020). WormCat: An Online Tool for Annotation and Visualization of *Caenorhabditis elegans* Genome-Scale Data. Genetics 214, 279–294.

15. Hwang, S., Lee, J., and Lee, S.V. (2025). Brief guide to assays for measuring health parameters using *Caenorhabditis elegans*. Mol Cells 48, 100233.

16. Jacobson, K.A., Gao, Z.G., Matricon, P., Eddy, M.T., and Carlsson, J. (2022). Adenosine A(2A) receptor antagonists: from caffeine to selective non-xanthines. Br J Pharmacol 179, 3496–3511.

17. Ji, Y., Jeon, Y.G., Lee, W.T., Han, J.S., Shin, K.C., Huh, J.Y., and Kim, J.B. (2024). PKA regulates autophagy through lipolysis during fasting. Mol Cells 47, 100149.

18. Jung, Y., Artan, M., Kim, N., Yeom, J., Hwang, A.B., Jeong, D.E., Altintas, Ö., Seo, K., Seo, M., Lee, D., et al. (2021). MON-2, a Golgi protein, mediates autophagy-dependent longevity in *Caenorhabditis elegans*. Sci Adv 7, eabj8156.

19. Kanehisa, M. and Goto, S. (2000). KEGG: kyoto encyclopedia of genes and genomes. Nucleic Acids Res 28, 27–30.

20. Kang, E., Kang, C., Lee, Y.S., and Lee, S.V. (2024). Brief guide to senescence assays using cultured mammalian cells. Mol Cells 47, 100102.

21. Kapahi, P., Kaeberlein, M., and Hansen, M. (2017). Dietary restriction and lifespan: Lessons from invertebrate models. Ageing Res Rev 39, 3–14.

22. Kim, D.Y., Moon, K.M., Heo, W., Du, E.J., Park, C.G., Cho, J., Hahm, J.H., Suh, B.C., Kang, K., and Kim, K. (2024). A FMRFamide-like neuropeptide FLP-12 signaling regulates head locomotive behaviors in *Caenorhabditis elegans*. Mol Cells 47, 100124.

23. Kim, E., Annibal, A., Lee, Y., Park, H.H., Ham, S., Jeong, D.E., Kim, Y., Park, S., Kwon, S., Jung, Y., et al. (2023). Mitochondrial aconitase suppresses immunity by modulating oxaloacetate and the mitochondrial unfolded protein response. Nat Commun 14, 3716.

24. Kim, T.Y. and Lee, B.D. (2025). Current therapeutic strategies in Parkinson’s disease: Future perspectives. Mol Cells 48, 100274.

25. Kwon, H.C. and Lee, S.V. (2025). Brief guide to *Caenorhabditis elegans* survival assays. Mol Cells 48, 100232.

26. Lee, D., An, S.W.A., Jung, Y., Yamaoka, Y., Ryu, Y., Goh, G.Y.S., Beigi, A., Yang, J.S., Jung, G.Y., Ma, D.K., et al. (2019). MDT-15/MED15 permits longevity at low temperature via enhancing lipidostasis and proteostasis. PLoS Biol 17, e3000415.

27. Lee, G.Y., Ham, S., and Lee, S.V. (2024a). Brief guide to RNA sequencing analysis for nonexperts in bioinformatics. Mol Cells 47, 100060.

28. Lee, G.Y., Ham, S., Sohn, J., Kwon, H.C., and Lee, S.V. (2024b). Meta-analysis of the transcriptome identifies aberrant RNA processing as common feature of aging in multiple species. Mol Cells 47, 100047.

29. Lee, J., Lee, B., Lee, H., Kim, E.J.E., Kim, S.S., Kwon, H.C., Lee, H., Lee, G.Y., Hong, W., Ham, S., et al. (2025a). Pelota-mediated ribosome-associated quality control counteracts aging and age-associated pathologies across species. Proc Natl Acad Sci U S A 122, e2505217122.

30. Lee, G.Y., Kim, S.J., Kwon, H.C., Sub, Y., Gee, H.Y., and Lee, S.V. (2025b). Functional Testing of Human Disease Missense Variants in *Caenorhabditis elegans* by Targeting COQ2 Variants. Kidney Int Rep 10, 3624–3639.

31. Lee, S.J., Lee, G.Y., Kim, S.S., Bae, Y., Ham, S., Sohn, J., Han, S.K., and Lee, S.V. (2025c). T-CLASS: An Online Tool for the Identification and Classification of Aging and Senescence Using Transcriptome Data. Aging Cell 24, e70193.

32. Lee, Y., Jung, Y., Jeong, D.E., Hwang, W., Ham, S., Park, H.H., Kwon, S., Ashraf, J.M., Murphy, C.T., and Lee, S.V. (2021). Reduced insulin/IGF1 signaling prevents immune aging via ZIP-10/bZIP-mediated feedforward loop. J Cell Biol 220.

33. Lopez-Otin, C., Blasco, M.A., Partridge, L., Serrano, M., and Kroemer, G. (2023). Hallmarks of aging: An expanding universe. Cell 186, 243–278.

34. McLellan, T.M., Caldwell, J.A., and Lieberman, H.R. (2016). A review of caffeine’s effects on cognitive, physical and occupational performance. Neurosci Biobehav Rev 71, 294–312.

35. Min, H., Kawasaki, I., Gong, J., and Shim, Y.H. (2015). Caffeine induces high expression of cyp-35A family genes and inhibits the early larval development in *Caenorhabditis elegans*. Mol Cells 38, 236–242.

36. Min, H., Park, G., and Lee, S.V. (2025). Brief guide to *Caenorhabditis elegans* imaging and quantification. Mol Cells 48, 100249.

37. Min, H., Youn, E., and Shim, Y.H. (2021). Long-Term Caffeine Intake Exerts Protective Effects on Intestinal Aging by Regulating Vitellogenesis and Mitochondrial Function in an Aged *Caenorhabditis elegans* Model. Nutrients 13.

38. Murphy, J.T., Liu, H., Ma, X., Shaver, A., Egan, B.M., Oh, C., Boyko, A., Mazer, T., Ang, S., Khopkar, R., et al. (2019). Simple nutrients bypass the requirement for HLH-30 in coupling lysosomal nutrient sensing to survival. PLoS Biol 17, e3000245.

39. Nehlig, A., Daval, J.L., and Debry, G. (1992). Caffeine and the central nervous system: mechanisms of action, biochemical, metabolic and psychostimulant effects. Brain Res Brain Res Rev 17, 139–170.

40. O’Rourke, E.J. and Ruvkun, G. (2013). MXL-3 and HLH-30 transcriptionally link lipolysis and autophagy to nutrient availability. Nat Cell Biol 15, 668–676.

41. Park, K., Jeon, M.C., Lee, D., Kim, J.I., and Im, S.W. (2024). Genetic and epigenetic alterations in aging and rejuvenation of human. Mol Cells 47, 100137.

42. Park, Y.J., Moon, K.M., and Kim, K. (2025). A practical guide to ordering *C. elegans* strains for biological research. Mol Cells 48, 100251.

43. Phung, O.J., Baker, W.L., Matthews, L.J., Lanosa, M., Thorne, A., and Coleman, C.I. (2010). Effect of green tea catechins with or without caffeine on anthropometric measures: a systematic review and meta-analysis. Am J Clin Nutr 91, 73–81.

44. Poole, R., Kennedy, O.J., Roderick, P., Fallowfield, J.A., Hayes, P.C., and Parkes, J. (2017). Coffee consumption and health: umbrella review of meta-analyses of multiple health outcomes. BMJ 359, j5024.

45. Reiner, Z., Guardamagna, O., Nair, D., Soran, H., Hovingh, K., Bertolini, S., Jones, S., Coric, M., Calandra, S., Hamilton, J., et al. (2014). Lysosomal acid lipase deficiency--an under-recognized cause of dyslipidaemia and liver dysfunction. Atherosclerosis 235, 21–30.

46. Ribeiro, J.A. and Sebastiao, A.M. (2010). Caffeine and adenosine. J Alzheimers Dis 20 Suppl 1, S3–15.

47. Santamarina-Fojo, S., Gonzalez-Navarro, H., Freeman, L., Wagner, E., and Nong, Z. (2004). Hepatic lipase, lipoprotein metabolism, and atherogenesis. Arterioscler Thromb Vasc Biol 24, 1750–1754.

48. Smith, A. (2002). Effects of caffeine on human behavior. Food Chem Toxicol 40, 1243–1255.

49. Sohn, J., Kwon, S., Lee, G.Y., Kim, S.S., Lee, Y., Lee, J., Jung, Y., Ham, S., Park, H.H., Park, S., et al. (2025). HLH-30/TFEB mediates sexual dimorphism in immunity in *Caenorhabditis elegans*. Autophagy 21, 283–297.

50. Spitler, K.M. and Davies, B.S.J. (2020). Aging and plasma triglyceride metabolism. J Lipid Res 61, 1161–1167.

51. Sutphin, G.L., Bishop, E., Yanos, M.E., Moller, R.M., and Kaeberlein, M. (2012). Caffeine extends life span, improves healthspan, and delays age-associated pathology in *Caenorhabditis elegans*. Longev Healthspan 1, 9.

52. Tabrez, S.S., Sharma, R.D., Jain, V., Siddiqui, A.A., and Mukhopadhyay, A. (2017). Differential alternative splicing coupled to nonsense-mediated decay of mRNA ensures dietary restriction-induced longevity. Nat Commun 8, 306.

53. Tabrizi, R., Saneei, P., Lankarani, K.B., Akbari, M., Kolahdooz, F., Esmaillzadeh, A., Nadi-Ravandi, S., Mazoochi, M., and Asemi, Z. (2019). The effects of caffeine intake on weight loss: a systematic review and dos-response meta-analysis of randomized controlled trials. Crit Rev Food Sci Nutr 59, 2688–2696.

54. Wanke, V., Cameroni, E., Uotila, A., Piccolis, M., Urban, J., Loewith, R., and De Virgilio, C. (2008). Caffeine extends yeast lifespan by targeting TORC1. Mol Microbiol 69, 277–285.

55. Yan, J., Chen, S., Yi, Z., Zhao, R., Zhu, J., Ding, S., and Wu, J. (2024). The role of p21 in cellular senescence and aging-related diseases. Mol Cells 47, 100113.

56. Yun, S.M., Park, J.Y., Seo, S.W., and Song, J. (2019). Association of plasma endothelial lipase levels on cognitive impairment. BMC Psychiatry 19, 187.

