## Supplementary Information for "Caffeine extends lifespan by enhancing lysosomal lipolysis in *Caenorhabditis elegans*"

**Supplementary Figures**

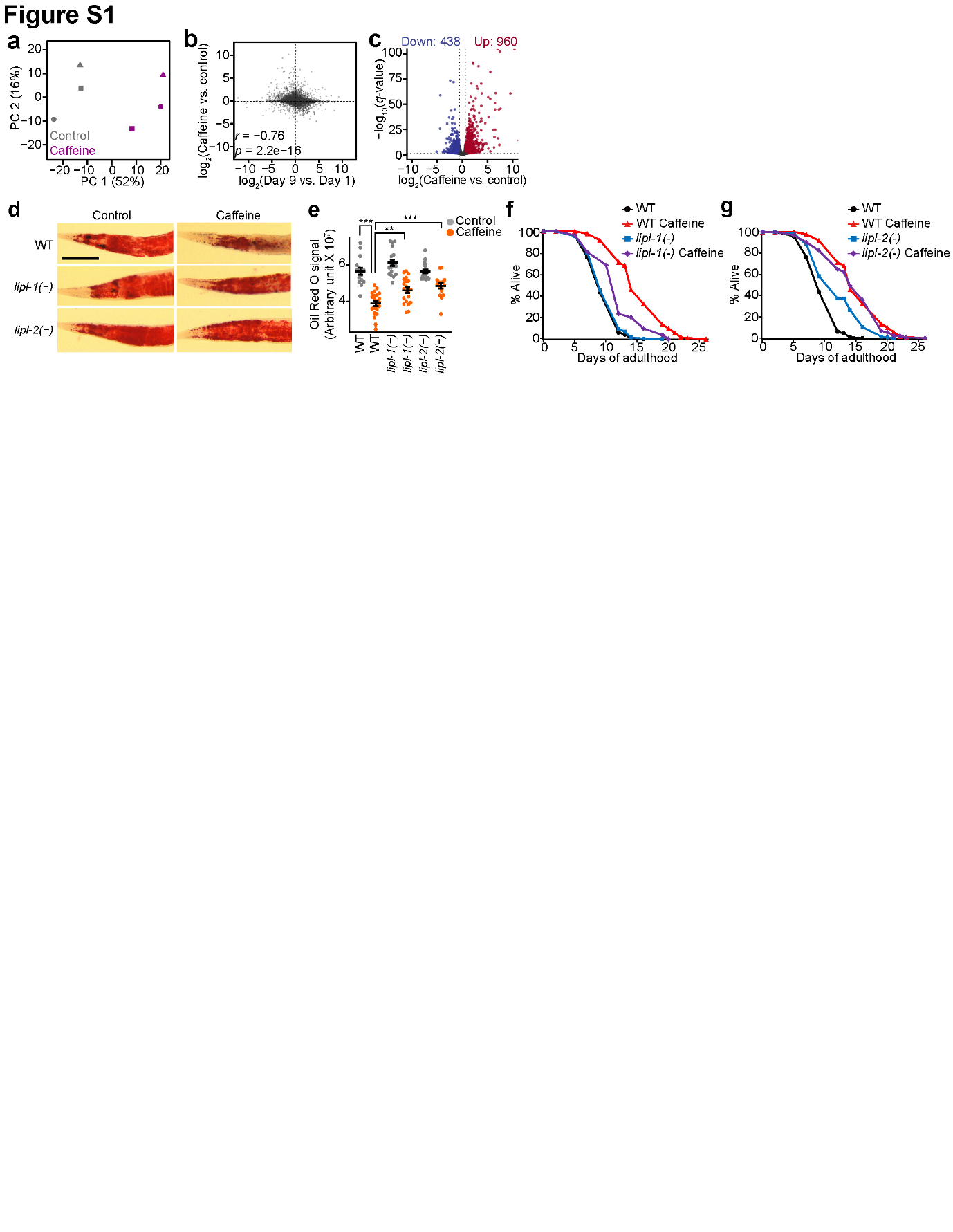

**Figure S1.** Caffeine-induced fat loss requires *lipl-1* and *lipl-2*, whereas lifespan extension is predominantly *lipl-1*-dependent. (**a**) Principal component analysis (PCA) plots of transcriptomic changes induced by caffeine feeding. PC, principal component. (**b**) A scatter plot comparing gene expression changes caused by caffeine feeding and by aging. Transcriptomic profiles of young (day 1 of adulthood) and old (day 9 of adulthood) animals were obtained by re-analyzing our previously published dataset (Lee et al., 2021). *r*, Pearson correlation coefficient. (**c**) Volcano plot showing differentially expressed genes by caffeine feeding. Genes with fold change > 1.5 and adjusted *p*-value < 0.05 were defined as differentially expressed genes. (**d**–**e**) Representative images (d) and quantification (e) of Oil Red O staining in wild-type (WT), *lipl-1(glm77)* [*lipl-1(−)*], and *lipl-2(**glm78)* [*lipl-1(−)*] mutants fed with or without caffeine (N ≥ 16 for each condition, two independent trials). Both *lipl-1(−)* and *lipl-2(−)* single mutations partially suppressed decreased lipid levels by caffeine feeding, with the suppression being slightly more pronounced by the *lipl-2(−)* mutations. To visualize the differential effects of caffeine on lipid storage in WT, *lipl-1(−)*, and *lipl-2(−)* single mutants, the same WT data sets are also shown in Fig. 1q and Fig. 1r. Scale bar, 75 μm. Error bars represent standard error of the mean (SEM). *p*-values were calculated by using one-way ANOVA (***p* < 0.01, ****p* < 0.001). (**f**–**g**) The lifespan of WT, *lipl-1(−)* (f), and *lipl-2(−)* (g) mutants fed with or without caffeine. *lipl-1* was partially required for the lifespan-extending effect of caffeine feeding (f), whereas *lipl-2(−)* did not affect the caffeine-induced longevity (g). To visualize the effects of caffeine on longevity in *lipl-1(−)* and *lipl-2(−)* single mutants, the same WT data sets are also shown in Fig. 1s. Lifespan results in this figure represent pooled data from two independent experiments; results of individual replicates are provided in Table S3.

**Supplementary Materials and Methods**

***C. elegans* strains**

*C. elegans* strains were maintained at 20°C on standard nematode growth medium (NGM) plates seeded with OP50 (Park et al., 2025). Strains used in this study are as follows: N2 wild-type (WT), IJ173 *eat-2(ad1116) II* (outcrossed at least four times to WT N2 strain), *CF2553 osm-5(p813) X* (outcrossed three times to WT N2 strain), AGP347 *lipl-1(glm77) V*, AGP364a *lipl-2(glm78) V*, AGP357a *lipl-2(glm79) lipl-1(glm77)* *V*, AGP334 *plipl-1:mCherry glmEx75[plipl-1(441bp)::mCherry + pmyo-3::GFP]*, AGP335 *plipl-2:mCherry glmEx76 [plipl-2(1.5kb)::mCherry + pofm-1::GFP]*

**RNA library preparation and sequencing**

WT animals were grown at 25°C and fed either with or without 10 mM caffeine for 72 h. Approximately 300 animals per biological replicate were harvested in TRIzol reagent and submitted to Macrogen Inc. (Seoul, Republic of Korea) for total RNA isolation, library preparation using TruSeq Stranded mRNA Library Prep Kit, and sequencing on the Illumina NovaSeq platform with paired-end reads.

**Analysis of RNA-seq data**

RNA sequencing data were analyzed following previous studies with minor modifications (Ham et al., 2022; 2024; Lee et al., 2024a; Lee et al., 2024b; Sohn et al., 2025). Sequencing reads were aligned to the *C. elegans* genome WBcel235 (ce11) and Ensembl transcriptome (release 104) by using STAR (v2.7.0e) (Dobin et al., 2013). Reads aligned to genes and transcripts were quantified by using RSEM (v.1.3.1) (Li and Dewey, 2011). Genes with fold change > 1.5 and adjusted *p* < 0.05 were defined as differentially expressed genes by caffeine supplementation by using DESeq2 (v1.28.0.) (Love et al., 2014). RNA-seq analyses of the published datasets PRJNA342407 (Tabrez et al., 2017) and GSE137861 (Lee et al., 2021) were performed using the same methods. Genes with fold change > 1.2 and adjusted *p* < 0.05 were defined as differentially expressed genes by *eat-2* mutations. Overrepresentation analysis of gene ontology (GO) cellular component (CC) terms and Kyoto Encyclopedia of Genes and Genomes (KEGG) (Kanehisa and Goto, 2000) was performed by using WebGestalt (Elizarraras et al., 2024). Transcriptomic CLassification via Adaptive learning of Signature States (T-CLASS) (Lee et al., 2025; http://www.t-class.kaist.ac.kr) was used to classify transcriptomic changes caused by caffeine into the three established longevity-promoting regimens in *C. elegans*.

**Lifespan assay using *C. elegans***

Lifespan assays using *C. elegans* were performed based on a previous study (Ji et al., 2024; Lee et al., 2025a; Kwon and Lee, 2025) with minor modifications. Streptomycin-resistant OP50 *Escherichia coli* bacteria were cultured in Luria broth (LB) media containing 10 μg/ml streptomycin at 37°C overnight. The cultured OP50 (100 μl) were seeded on NGM with or without 10 mM caffeine and incubated at 37°C overnight. To prevent progeny from hatching, the DNA synthesis inhibitor 5-fluoro-2′-deoxyuridine (FUDR) (Sigma-Aldrich, F0503, USA) was added to the OP50-seeded plates at a final concentration of 5 μM. Animals were cultured from egg to L4 larval stage on OP50-seeded NGM plates at 25°C, transferred to freshly prepared plates with or without 10 mM caffeine, and then transferred to new plates after one or two days. Lifespan assays were conducted at 25°C on NGM plates. *C. elegans* that did not respond to a gentle touch using a platinum wire were counted as dead. Animals that displayed rupture or internal hatching, burrowed, or crawled off the plates were censored but included in statistical analysis. Online application of survival analysis 2 (OASIS2, http://sbi.postech.ac.kr/oasis2) (Han et al., 2016) and OASIS portable (OASISp) (Han et al., 2024) were used for statistical analysis of lifespan data. The log-rank (Mantel-Cox method) test was used to calculate *p*-values.

**Microscopy and image analysis**

For microscopy, L4 larval stage animals were transferred to OP50-seeded NGM plates with or without 10 mM caffeine and maintained at 25°C. Fluorescence images of *plipl-1::mCherry-* and *plipl-2::mCherry*-expressing animals (Bahr et al., 2025) were obtained after 72 h on these plates. Animals were then mounted on a 2% agarose pad and anesthetized with 100 mM sodium azide (Sigma-Aldrich, 71289-5 G) prior to imaging (Jung et al., 2021; Kim et al., 2024; Min et al., 2025). Images were acquired using a Leica DFC9000 GT deep-cooled sCMOS camera and Leica LAS X imaging software (Leica Microsystems, Germany). ImageJ was used to quantify the fluorescence intensity (Schneider et al., 2012; Mehlem et al., 2013), and the background signals were subtracted.

**Measurement of pharyngeal pumping rates**

Pharyngeal pumping (feeding) of *C. elegans* was measured as previously described (Hwang et al., 2025; Lee et al., 2025b) with minor modifications. Animals were cultured from egg to L4 larval stage on OP50-seeded NGM plates at 20°C and then transferred to NGM plates with or without 10 mM caffeine. Animals were maintained on these plates for 72 h at 25°C. The pharyngeal pumping rate was counted for 30 seconds and then converted to the number of pumps per minute. The rates of pumping were observed under a dissecting microscope (Zeiss SteREO Discovery V8, Zeiss Corporation, Jena, Germany). Each assay was performed in two independent experiments. Two-tailed *t*-test was used for statistical analysis.

**RNA extraction and quantitative RT-PCR (RT-qPCR)**

RNA extraction and RT-qPCR were performed as previously described (Kim et al., 2023; Bong et al., 2024) with minor modifications. Synchronized L4 larval stage animals were transferred to NGM plates with or without 10 mM caffeine and maintained for 72 h at 25°C. Animals were harvested and washed three times with M9 buffer containing 0.01% polyethylene glycol 4000 (PEG4000, Tokyo, Chemical Industry, Tokyo, Japan). Total RNA was extracted using RNAiso plus (Takara, Seta, Kyoto, Japan) and cDNA template was synthesized using ImProm-II™ Reverse Transcriptase kit (Promega, Madison, WI, USA) with random primers. StepOne Real-Time PCR System (Applied Biosystems, Foster City, CA, USA) and SYBR Green Master Mix (4367659, Applied Biosystems, Foster City, CA, USA) were utilized for performing RT-qPCR following the manufacturer’s protocol. Comparative C_T_ method was used for the quantitative analysis of mRNAs. The mRNA level of *tba-1*, which encodes a tubulin α, was used as a normalization control. Primer sequences used for RT-qPCR analyses in this study are listed in Table S1.

**Oil Red O staining**

Oil Red O staining was performed as previously described (Lee et al., 2019) with minor modifications. Synchronized L4 larval stage animals were cultured on OP50-seeded NGM plates with or without 10 mM caffeine for 72 h at 25°C. Animals were then harvested in M9 buffer, washed three times with PBST (137 mM NaCl, 2.7 mM KCl, 10 mM Na_2_HPO_4_, 2 mM KH_2_PO_4_ with 0.01% Triton X-100), and fixed with 50% isopropanol for 3 min. Oil Red O solution (0.5% in isopropanol, Sigma-Aldrich) was diluted with double-distilled water (ddH_2_O) to 60% to prepare the working solution and precipitates were removed by filtering. Fixed animals were incubated in the working solution for 2 h at room temperature. Stained animals were washed with PBST and mounted on 2% agarose pads. Differential interference contrast (DIC) images were acquired using a Leica DFC9000 GT deep-cooled sCMOS camera and Leica LAS X imaging software (Leica Microsystems, Germany). ImageJ was used for the quantification of Oil Red O intensity (Schneider et al., 2012; Mehlem et al., 2013). For image analysis, images were converted to 8-bit format and processed using the “Invert LUTs” function in ImageJ. For quantification of Oil Red O staining, a fixed threshold value of 120 was applied, and areas with intensities above this threshold in the anterior intestinal cells were measured.

**Supplementary Information References**

1. Bahr, L., Amrit, F.R., Silvia, P.E., Wayhs, B., Osman, G.A., Devare, M.N., Henry, H., Bui, D., Choe, M., and Naim, N. (2025). LIPL-1 and LIPL-2 are TCER-1-regulated Lysosomal Lipases with Distinct Roles in Immunity and Fertility. PLoS genetics 21, e1011804.
2. Bong, D., Sohn, J., and Lee, S.V. (2024). Brief guide to RT-qPCR. Mol Cells 47, 100141.
3. Dobin, A., Davis, C.A., Schlesinger, F., Drenkow, J., Zaleski, C., Jha, S., Batut, P., Chaisson, M., and Gingeras, T.R. (2013). STAR: ultrafast universal RNA-seq aligner. Bioinformatics 29, 15-21.
4. Elizarraras, J.M., Liao, Y., Shi, Z., Zhu, Q., Pico, A.R., and Zhang, B. (2024). WebGestalt 2024: faster gene set analysis and new support for metabolomics and multi-omics. Nucleic Acids Res 52, W415-W421.
5. Ham, S., Kim, S.S., Park, S., Kim, E.J.E., Kwon, S., Park, H.H., Jung, Y., and Lee, S.V. (2022). Systematic transcriptome analysis associated with physiological and chronological aging in *Caenorhabditis elegans*. Genome Res 32, 2003-2014.
6. Ham, S., Kim, S.S., Park, S., Kwon, H.C., Ha, S.G., Bae, Y., Lee, G.Y., and Lee, S.V. (2024). Combinatorial transcriptomic and genetic dissection of insulin/IGF-1 signaling-regulated longevity in *Caenorhabditis elegans*. Aging Cell 23, e14151.
7. Han, S.K., Kwon, H.C., Yang, J.S., Kim, S., and Lee, S.V. (2024). OASIS portable: User-friendly offline suite for secure survival analysis. Mol Cells 47, 100011.
8. Han, S.K., Lee, D., Lee, H., Kim, D., Son, H.G., Yang, J.S., Lee, S.V., and Kim, S. (2016). OASIS 2: online application for survival analysis 2 with features for the analysis of maximal lifespan and healthspan in aging research. Oncotarget 7, 56147-56152.
9. Hwang, S., Lee, J., and Lee, S.V. (2025). Brief guide to assays for measuring health parameters using *Caenorhabditis elegans*. Mol Cells 48, 100233.
10. Ji, Y., Jeon, Y.G., Lee, W.T., Han, J.S., Shin, K.C., Huh, J.Y., and Kim, J.B. (2024). PKA regulates autophagy through lipolysis during fasting. Mol Cells 47, 100149.
11. Jung, Y., Artan, M., Kim, N., Yeom, J., Hwang, A.B., Jeong, D.E., Altintas, Ö., Seo, K., Seo, M., Lee, D., et al. (2021). MON-2, a Golgi protein, mediates autophagy-dependent longevity in *Caenorhabditis elegans*. Sci Adv 7, eabj8156.
12. Kanehisa, M. and Goto, S. (2000). KEGG: kyoto encyclopedia of genes and genomes. Nucleic Acids Res 28, 27-30.
13. Kim, D.Y., Moon, K.M., Heo, W., Du, E.J., Park, C.G., Cho, J., Hahm, J.H., Suh, B.C., Kang, K., and Kim, K. (2024). A FMRFamide-like neuropeptide FLP-12 signaling regulates head locomotive behaviors in *Caenorhabditis elegans*. Mol Cells 47, 100124.
14. Kim, E., Annibal, A., Lee, Y., Park, H.H., Ham, S., Jeong, D.E., Kim, Y., Park, S., Kwon, S., Jung, Y., et al. (2023). Mitochondrial aconitase suppresses immunity by modulating oxaloacetate and the mitochondrial unfolded protein response. Nat Commun 14, 3716.
15. Kwon, H.C. and Lee, S.V. (2025). Brief guide to *Caenorhabditis elegans* survival assays. Mol Cells 48, 100232.
16. Lee, D., An, S.W.A., Jung, Y., Yamaoka, Y., Ryu, Y., Goh, G.Y.S., Beigi, A., Yang, J.S., Jung, G.Y., Ma, D.K., et al. (2019). MDT-15/MED15 permits longevity at low temperature via enhancing lipidostasis and proteostasis. PLoS Biol 17, e3000415.
17. Lee, G.Y., Ham, S., and Lee, S.V. (2024a). Brief guide to RNA sequencing analysis for nonexperts in bioinformatics. Mol Cells 47, 100060.
18. Lee, G.Y., Ham, S., Sohn, J., Kwon, H.C., and Lee, S.V. (2024b). Meta-analysis of the transcriptome identifies aberrant RNA processing as common feature of aging in multiple species. Mol Cells 47, 100047.
19. Lee, J., Lee, B., Lee, H., Kim, E.J.E., Kim, S.S., Kwon, H.C., Lee, H., Lee, G.Y., Hong, W., Ham, S., et al. (2025a). Pelota-mediated ribosome-associated quality control counteracts aging and age-associated pathologies across species. Proc Natl Acad Sci U S A 122, e2505217122.
20. Lee, G.Y., Kim, S.J., Kwon, H.C., Sub, Y., Gee, H.Y., and Lee, S.V. (2025b). Functional Testing of Human Disease Missense Variants in *Caenorhabditis elegans* by Targeting COQ2 Variants. Kidney Int Rep 10, 3624-3639.
21. Lee, S.J., Lee, G.Y., Kim, S.S., Bae, Y., Ham, S., Sohn, J., Han, S.K., and Lee, S.V. (2025c). T-CLASS: An Online Tool for the Identification and Classification of Aging and Senescence Using Transcriptome Data. Aging Cell 24, e70193.
22. Lee, Y., Jung, Y., Jeong, D.E., Hwang, W., Ham, S., Park, H.H., Kwon, S., Ashraf, J.M., Murphy, C.T., and Lee, S.V. (2021). Reduced insulin/IGF1 signaling prevents immune aging via ZIP-10/bZIP-mediated feedforward loop. J Cell Biol 220.
23. Li, B. and Dewey, C.N. (2011). RSEM: accurate transcript quantification from RNA-Seq data with or without a reference genome. BMC Bioinformatics 12, 323.
24. Love, M.I., Huber, W., and Anders, S. (2014). Moderated estimation of fold change and dispersion for RNA-seq data with DESeq2. Genome Biol 15, 550.
25. Mehlem, A., Hagberg, C.E., Muhl, L., Eriksson, U., and Falkevall, A. (2013). Imaging of neutral lipids by oil red O for analyzing the metabolic status in health and disease. Nat Protoc 8, 1149-1154.
26. Min, H., Park, G., and Lee, S.V. (2025). Brief guide to *Caenorhabditis elegans* imaging and quantification. Mol Cells 48, 100249.
27. Park, Y.J., Moon, K.M., and Kim, K. (2025). A practical guide to ordering C. elegans strains for biological research. Mol Cells 48, 100251.
28. Schneider, C.A., Rasband, W.S., and Eliceiri, K.W. (2012). NIH Image to ImageJ: 25 years of image analysis. Nat Methods 9, 671-675.
29. Sohn, J., Kwon, S., Lee, G.Y., Kim, S.S., Lee, Y., Lee, J., Jung, Y., Ham, S., Park, H.H., Park, S., et al. (2025). HLH-30/TFEB mediates sexual dimorphism in immunity in *Caenorhabditis elegans*. Autophagy 21, 283-297.
30. Tabrez, S.S., Sharma, R.D., Jain, V., Siddiqui, A.A., and Mukhopadhyay, A. (2017). Differential alternative splicing coupled to nonsense-mediated decay of mRNA ensures dietary restriction-induced longevity. Nat Commun 8, 306.
